# Proactive Versus Reactive Control Strategies Differentially Mediate Alcohol Drinking in Wistar and P rats

**DOI:** 10.1101/2023.06.08.544260

**Authors:** M.D. Morningstar, N.M. Timme, B. Ma, E. Cornwell, T. Galbari, C.C. Lapish

## Abstract

Problematic alcohol consumption is associated with deficits in decision-making, and alterations in prefrontal cortex neural activity likely contributes. We hypothesized that differences in cognitive control would be evident between male Wistar rats and a model for genetic risk for alcohol use disorder (alcohol-preferring P rats). Cognitive control can be split into proactive and reactive components. Proactive control maintains goal-directed behavior independent of a stimulus whereas reactive control elicits goal-directed behavior at the time of a stimulus. We hypothesized that Wistars would show proactive control over alcohol-seeking whereas P rats would show reactive control over alcohol-seeking. Neural ensembles were recorded from prefrontal cortex during an alcohol seeking task that utilized two session types. On congruent sessions the CS+ was on the same side as alcohol access. Incongruent sessions presented alcohol opposite the CS+. Wistars, but not P rats, exhibited an increase in incorrect approaches during incongruent sessions, suggesting that Wistars utilized the previously learned task-rule. This motivated the hypothesis that ensemble activity reflecting proactive control would be observable in Wistars but not P rats. While P rats showed differences in neural activity at times relevant for alcohol delivery, Wistars showed differences prior to approaching the sipper. These results support our hypothesis that Wistars are more likely to engage proactive cognitive-control strategies whereas P rats are more likely to engage reactive cognitive control strategies. Although P rats were bred to prefer alcohol, differences in cognitive control may reflect a sequela of behaviors that mirror those in humans at risk for an AUD.

**Significance Statement:** Cognitive control refers to the set of executive functions necessary for goal-directed behavior. It is a major mediator of addictive behaviors and can be subdivided into proactive and reactive cognitive control. We observed behavioral and electrophysiological differences between outbred Wistar rats and the selectively bred Indiana alcohol-preferring P rat while they sought and consumed alcohol. These differences are best explained by reactive cognitive control in P rats and proactive in Wistar rats.

## Introduction

Understanding how interactions between alcohol history and genotype impact cognitive function is critical for elucidating the mechanisms behind alcohol use disorder (AUD) (Wilcox et al., 2014). We sought to determine if differences between alcohol seeking in P rats and Wistars could be explained as differences in cognitive control strategies. Cognitive control is the set of executive functions necessary for reducing uncertainty prior to an action (Mackie et al., 2013), and can be broken into dual mechanisms of proactive and reactive control (Braver, 2012). Proactive cognitive control maintains an active representation of the goal at hand; reactive control draws upon the representation when elicited by a cue or stimulus (Braver, 2012). Cognitive control is linked with AUD both as a risk factor as well as a domain that AUD disrupts (Wilcox et al., 2014; Liu et al., 2020). Refining our understanding of how subtypes of cognitive control may interact with AUD is critical. We investigated the degree to which P rats and Wistar rats differentially instantiate cognitive control and hypothesized that Wistars are more likely to implement proactive cognitive control, whereas P rats are more likely to implement reactive cognitive control.

The 2-way conditioned access protocol (2CAP) has been previously used to assess how heritable factors result in differences in neural activity and alcohol seeking behavior. (Bell et al., 2006; McCane et. al, 2014; Linsenbardt & Lapish, 2015, Linsenbardt et al., 2019, Timme et al., 2020, Timme et al., 2022). P rats display high alcohol seeking and consumption behaviors (Czachowski & Samson, 2002). P rats also display high levels of impulsivity (Beckwith & Czachowski, 2014; Linsenbardt et al., 2017) and impaired cognitive flexibility (De Falco et al., 2021). Increased impulsivity and decreased cognitive flexibility may be a result of decreased representations of task goals. For example, previous work has consistently shown blunted neural activity in dorsal-medial prefrontal cortex (dmPFC) prior to alcohol seeking in P, but not Wistar, rats (Linsenbardt & Lapish, 2015; Linsenbardt et al., 2019; Timme et al., 2022).

The dmPFC is critical for cognitive control (Braver, 2012). A general working model of dmPFC function is to test for discrepancies, and if found, modulate attentional or emotive resources to address the discrepancy (Alexander & Brown, 2011). The dmPFC must maintain a variety of stimulus-response-outcomes whose representations can be thought of as task sets (Wisniewski et al., 2015; de Haan et al., 2018; Soltani & Koechlin, 2022). Task sets are neural representations critical for the rapid updating of learned behavior and assist in proactive control (Dreisbach & Haider, 2008; Dhawan et al., 2019; Soltani & Koechlin, 2022). Therefore, it would be expected that agents that engage proactive control would exhibit strong task-sets.

An agent that is more reactive in their alcohol seeking is not required to maintain a goal representation, and goal representations may emerge only when needed. In the 2CAP task this would correspond to periods of alcohol access. Reactive control is correlated with impulsivity (Huang et al., 2017) and impulsivity is predictive of compulsive drinking (Finn et al., 1999; Acheson et al., 2011), therefore it is expected that reactive control is also predicted of compulsive drinking. This may partially explain why P rats are more likely to drink through quinine adulterated alcohol (Timme et al., 2022). High yield *in vivo* electrophysiology will allow us to determine if changes in ensembles activity reflect differences in cognitive control between P’s and Wistars.

We examined the differences between low-drinking Wistars and high-drinking P rats as they adapted to a change in contingency during the alcohol seeking 2CAP task. Given that P rats are more impulsive and less flexible, we hypothesized that they would utilize reactive cognitive control strategies that do not depend upon maintained goal representations.

Conversely, we hypothesized our Wistar population would utilize proactive cognitive control strategies. The hypothesized differences in cognitive control would be evident in both the behavioral and electrophysiological data.

## MATERIALS AND METHODS

### Data

The present dataset was produced concurrently with Timme et al., 2022. Previous papers analyzed the effects of quinine on congruent (regular) 2CAP sessions. The present paper utilizes unpublished data from previously unexplored congruent and incongruent sessions. Congruent sessions are obtained the day prior to incongruent sessions.

### Animals

Adult, male Indiana Alcohol Preferring rats (P rats) were utilized from breeding facilities at the Indiana University School of Medicine. Adult, male Wistar rats were acquired from Envigo (Envigo, IN). 60 Wistars and 23 P rats were allowed to drink in an intermittent access protocol (IAP). Following IAP, 20 Wistars and 16 P rats were selected for further 2CAP training. From those 20 Wistars and 16 P rats, 8 Wistars and 8 P rats were implanted with silicon electrodes. One Wistar had no incongruent datasets. In total, 7 Wistars and 8 P rats were used in the present dataset. Animals were kept on a reversed light schedule and given standard rat chow and water ad libitum. All animals were single housed after arriving in our colony and throughout the experiment. All animal procedures were approved by the Indiana University – Purdue University Indianapolis School of Science Institutional Animal Care and Use Committee.

### Intermittent access protocol

All animals underwent an intermittent access protocol (IAP) wherein they were given periodic access to 20% v/v alcohol for a 2-week period. Specifically, animals were weighed and then a bottle of water and a bottle of alcohol were attached to each animals’ cage for a period of 24 hours. Following those 24 hours, the water bottle was weighed and the alcohol bottle was weighed. The difference between the initial and final weight was obtained and from this the dose consumed was calculated by dividing this difference by the animal’s weight (g/kg). Bottles were placed and animals were weighed specifically on Mondays, Wednesdays, and Fridays 2-hours into the animals’ dark cycle. Following IAP, high-drinking Wistars and lower-drinking P rats were selected to go on to 2CAP training (Timme et al., 2022).

### Apparatus

Animals underwent 2CAP training in a standard Med Associates shuttle box (Med Associates, VT). All electrophysiology recordings were completed in a replica Med Associates shuttle box that allowed passage of our tethers and RGB video tracking.

### 2-way cued access protocol

Animals underwent training in the 2-way cued access protocol (2CAP) for a period of 2 weeks. 2CAP training consisted of presentations of a conditioned stimulus (CS+) prior to the presentation of a 10% v/v alcohol sipper. An additional, separate stimulus was presented (CS-) on trials where no alcohol sipper would descend. During CS+ trials, the CS+ would illuminate on a single side of the apparatus. During normal, congruent sessions, this CS+ matched the side of the apparatus where alcohol would be present. The CS-, in contrast, was illuminated on both sides of the 2CAP apparatus. **Figure 1** describes the contingency swap as well as apparatus.

**Figure 1.**
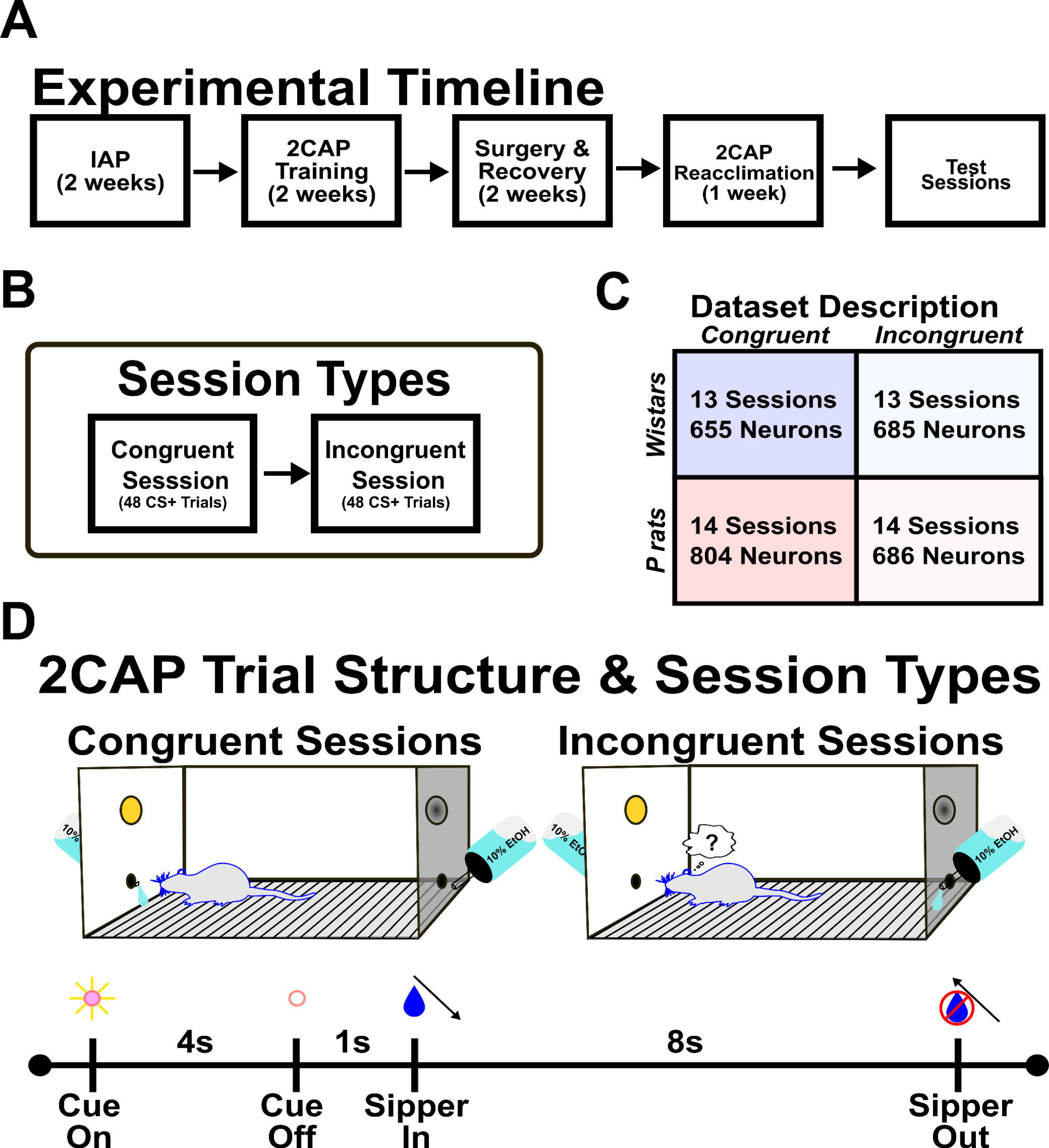
Method details and 2CAP apparatus. A. Wistars and P rats underwent 2 weeks of intermittent access protocol (IAP) with 20% alcohol (v/v). Following IAP, Wistars and P rats were matched for consumption and underwent subsequent 2CAP training with 10% alcohol (v/v). Following successful completion of 2CAP training, animals underwent surgery and had silicon electrodes chronically implanted. Animals were then reacclimated to the 2CAP protocol as well as habituated to the modified 2CAP chamber. **B.** After reacclimating, neural recordings were acquired. Each week progressed through different contingencies. We focused the present analysis on congruent sessions and incongruent sessions conducted at the end of the week. Each session type consisted of 48 CS+ trials and 48 CS- trials. **C.** Wistars and P rats experienced roughly equal numbers of congruent and incongruent sessions. Additionally, neuron yield was relatively matched. **D.** The 2CAP apparatus and trial structure is detailed. Briefly, during congruent sessions alcohol access was on the same side as the CS+. During incongruent sessions, alcohol access was on the side opposite the CS+. A 2CAP trial consisted of 4-seconds of the CS+ being illuminated. 1-second after the CS+ is turned off, the alcohol containing sipper would descend. Animals then had 8-seconds of 10% alcohol access prior to the sipper being removed from the chamber.

Both the CS+ and CS- were visual and could either be a solid light presented for 4 seconds or a blinking (1 Hz) light presented over the course of 4 seconds. The type of visual stimulus was assigned randomly and counter balanced. A single 2CAP session had 96 trials wherein 48 presented the CS+ on a random side of the apparatus and the other 48 trials presented the CS- on both sides of the apparatus. **Figure 1D** highlights the events of a single trial. During the sipper descent, both motors were active such that neither motor noise could be used as a discriminative stimulus. Following training and during electrophysiology recordings, animals were exposed to 1-3 incongruent sessions. Incongruent sessions were when the CS+ that normally predicted the side of alcohol delivery was swapped to predict the side absent alcohol. Thus, animals were required to adapt over a single session to the new contingency.

### Behavioral data

Behavioral data were processed with DeepLabCut (DLC) (Mathis et al., 2018). Video recordings were saved in color and at 30 FPS. The training dataset identified six points in the 2CAP apparatus (each corner, both sippers) and 4 points on the animals’ body (snout, head cap, back, tail). Following successful training of a neural network, all videos relevant to the current study were processed. Output data were a set of {X, Y} coordinates per identified point of interest along with the measure of confidence the neural network had for those {X, Y} coordinates at each time point. Data were preprocessed in MATLAB by interpolating timepoints that the neural network had low confidence in (<90% log-likelihood). This ultimately allowed for an estimate of the animals’ position at each time point relative to the correct or incorrect sippers. This then permitted quantification of correct versus incorrect sipper approaches, time spent at each sipper, and other metrics related to approach and drinking behavior with great temporal precision.

### Surgeries

Surgeries were performed on animals following IAP and 2CAP. Pre-surgery 2CAP drinking was considered in the selection of which animals would move forward in the experiments such that lower-drinking P rats were matched with higher-drinking Wistar rats. Briefly, animals were rendered unconscious under isoflurane and fur on their skull was removed. Animals were then placed in the stereotaxic frame. Animals received doses of ketoprofen (5 mg/kg, i.p.) and cefazolin (30 mg/kg, i.p.). The scalp surface was sanitized with alternating applications of betadine and 70% alcohol. Anesthetic depth was confirmed via checking muscle reflexes. Following sufficient depth of anesthesia, a topical analgesic was applied to the scalp surface (bupivacaine, 5 mg, s.c.), and then an incision was made revealing the skull. The skull surface was cleaned and cleared of all debris and tissue until bregma and lambda were clearly visible. Four holes were drilled for anchoring screws, two holes were drilled for grounding screws, and one craniotomy was performed at the site of probe placement (3.2 mm A/P, 0.8 mm M/L, 3.0 mm D/V, 7.5 degrees toward midline). The electrode itself was mounted on a movable microdrive that allowed for post-surgery adjustments to be made to the depth of the electrode. All portions of the microdrive and electrode that required mobility were coated in warmed antibiotic ointment prior to the application of the dental cement head cap.

Following construction of the headcap, all open connectors were taped shut and the animal was removed from the stereotaxic equipment and placed into a recovery cage located atop a heating pad. The animal was monitored until they regained their righting reflex and then placed back into the vivarium. Postoperative health checks were done up to 7 days following the surgery.

### Electrophysiology

Implanted silicon electrodes were purchased from Cambridge Neurotech (Cambridge Neurotech, UK) and consisted of 3 different models: P, F, and H with most electrodes being of the F model. The only difference between models is the geometry of the recording sites. All electrodes came precoated in PEDOT allowing for greater biocompatibility. Omnetic (Omnetics Connector Corporation, MN) connectors and Intan (Intan, CA) 32 channel or 64 channel headstage pre-amplifiers were utilized in all recordings. Intan SPI cables were utilized to connect the headstages to our OpenEphys acquisition board (OpenEphys, GA). Raw electrophysiological data were sampled and recorded at 30,000 Hz via an OpenEphys acquisition board. Spikesorting was done offline via Kilosort 2.0 (Pachitariu et al., 2016).

Putative units were manually separated into noise, multi-unit activity, or single-unit activity via Phy (https://github.com/cortex-lab/phy). From there, only single units that had fewer than 5% inter-spike interval violations were considered in further analyses. Electrode placements can be found in Timme et al., 2022.

### Statistical analysis

Data were assessed for normality in all cases and either parametric or non-parametric tests were utilized afterwards. All statistics were done in MATLAB 2021a (The Mathworks, MA) using customized scripts or JASP using exported CSV files from MATLAB. Multiple comparisons were conducted where appropriate and unless otherwise stated were FDR corrected.

### Data analysis and software

#### General information

MATLAB 2021b and MATLAB 2021a were utilized to operate Kilosort 2.0 as well as process all subsequent behavioral and electrophysiological data. Python 3 was utilized to execute DLC scripts and the Phy GUI. Custom MATLAB scripts were utilized for all data processing and analysis.

#### Behavioral analyses

Data obtained and interpolated from DLC was utilized to determine the position of the animal at a temporal resolution of 0.033 ms. This allowed for precise quantification of the animals’ behaviors during each trial. Trials were either centered around the cue-onset or the time of correct approach. An approach was defined as the first timepoint when the DLC marker associated with the animals’ snout was less than or equal to 9 pixels in distance from the DLC marker associated with the sipper port. This information allowed us to quantify latencies, correct approaches, incorrect approaches, omissions, and so forth. Correct approaches were defined as trials where the animal immediately goes to the correct, alcohol-containing sipper without checking the other port. Incorrect approaches were defined as trials where the animal visited the port where alcohol was not made available. Omissions were when the animal visited neither port during a trial. Latencies were defined as the first timepoint the animal checked the port following the sipper coming into the arena. Behavioral data were selected [-5 20] seconds from the cue onset. The probability to approach any sipper was first encoded as either a 1 or 0. From there, the probabilities were split into correct or incorrect trials. A 3-trial moving average was calculated across each animals’ row of data to obtain a smoothed average. The probability to occupy the sipper was also calculated. In this analysis, a time-corresponding vector was loaded with either 1 or 0 depending on whether the animal was within 9 pixels of the sipper. Data were subsequently split into the occupancy of correct or incorrect sipper. A simple mean was then calculated across sessions to determine the time by trial average.

#### Firing rate

Data from the results of spikesorting was imported into MATLAB and transformed into a spike-timing matrix. Each column contained the spike times of an individual neuron and each row contained the Nth spike time. Firing rates were first binned at 100 ms. For each neuron a gaussian kernel was applied that considered the neurons’ average inter-spike interval (ISI) and its coefficient of variation (CV) in order to set the width of the smoothing kernel on an individual basis (Lehky, 2010). The width of each neuron’s individual smoothing kernel was determined by multiplying the square root of that neuron’s mean ISI with the inverse CV.

#### Electrophysiology analyses: centered on cue onset

The first set of analyses were concerned with the neural population signals relative to the cue onset. The time window used in these analyses was from 4 seconds prior to the cue until 18 seconds after the cue [-4 18] resulting in a total of 22 seconds of data. This time bin was adjusted from the behavioral analyses to minimize non-task related neural activity. Due to animals decreased responding over trials, only the first 15 trials were used in this analysis. This resulted in a 3-dimensional data structure with trials, time bins, and all neurons across their respective sessions, separated by strain. The mean over trials was taken for each session, and then data were concatenated. Principal component analysis (PCA) was then conducted with congruent and incongruent Wistar sessions undergoing PCA separately from congruent and incongruent P rat sessions. **Figure 5** describes the data structure prior to PCA. A broken stick model was utilized to determine how many PCs to include in any subsequent analyses (Barton & David, 1955; Frontier, 1976). Briefly, each component had a theoretical variance that it must exceed to be considered of relevance. The theoretical variance was calculated using Equation 1. The variance associated with each component is *b_k_*. Each component is one of p total. The variable k corresponds to the kth component analyzed. For example, when p = 7 the first component (k = 1) would need to exceed 37% of the explained variance to retain.

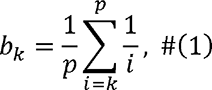

PC projections were calculated from the mean-centered data and coefficient loadings of each condition (Wistar congruent, Wistar incongruent, P rat congruent, P rat incongruent).

#### Electrophysiology analyses: centered on approach onset

The second set of analyses were primarily concerned with the activity 2 seconds before and 2 seconds after the animals’ approach to a sipper [-2 2]. The approach time was when the animals’ snout was less than or equal to 9 pixels from the sipper port during the period of fluid access. The time window was set to allow testing of whether strategies are encoded and implemented in a proactive manner with minimal interference from other task variables. The distance metrics for the approach were initially tested with the same PCA setup as described above. Following this, we sought to identify additional differences between approaches. We utilized the spatial nature of the 2CAP shuttle box to determine if distinct neural correlates existed for approaches to the left or right side of the box. To best compare how each strain encodes left and right approaches, a different PCA strategy was utilized. Briefly, we grouped data by the session type rather than strain. In this analysis, a mean calculated over trials was calculated for each session that contained up to 15 correct approaches for each direction (left or right). The mean of those session means was then calculated for each condition (Wistar congruent, Wistar incongruent, P rat congruent, P rat incongruent). This resulted in left and right means for each strain and session type. Prior to PCA, the means were concatenated in time such that the first half were the timepoints associated with the left trials and the second half were the timepoints associated with the right trials. **Figure 5** details the PCA strategies used for each analysis. Following PCA, data were sorted into left and right trials. First, we utilized a PCA permutation approach. Briefly, based on the broken stick model a set of permutations was generated using Equation 2 where n corresponds to the broken stick point and k corresponds to the desired amount of PC dimensions, which in our case was 3. 3 PC dimensions were utilized for computational simplicity, it is more intuitive, and it allows for ease of comparisons across session and strain types.

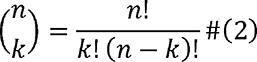

In each permutation, 1000 randomly selected left and right trials were compared in the 3D space. Their distance was taken at each sample. The mean distance for each permutation was then utilized as a general measure of separation for left and right trials in that strain or session type condition. Next a separate analysis was conducted where we analyzed raw firing rates.

Neurons that were less than or greater than 0.01 or -0.01 were sorted into their relevant coefficient loading. These values gave us roughly equal proportions of positive and negative loading neurons as well as a middle, undefined proportion that was not analyzed. The raw firing rate was calculated according to each PC’s positive and negative loaders. A repeated-measures ANOVA was utilized to find time by direction interactions within the PC population. PC4 was then chosen based on both the quantitative criteria and qualitative similarity across both congruent and incongruent sessions. The mean value before and after making the approach was then calculated for each session and strain combination and those values were compared. Last, the mean session latencies were compared with the mean distance values between left and right choices for each session. A linear regression and Pearson’s correlation was calculated.

## RESULTS

### Wistars are more likely to incorrectly approach the sipper during incongruent sessions

Wistars and P rats drank significantly different amounts throughout the task but did not show differences between session types. A main effect of strain was detected for the total alcohol intake (ANOVA, F(1,50) = 58.8, p < 0.001, **Figure 2A**). We additionally analyzed the total time at the sipper, and no main effect of strain (ANOVA, F(1,50) = 0.093, p = 0.762) or session type was detected (F(1,50) = 0.028, p = 0.866, **Figure 2B**). A main effect of strain (F(1,50) = 18.424, p < 0.001, **Figure 2C**) was found for the rate of intake. Despite the changed contingency, no session type differences in drinking were observed.

**Figure 2.**
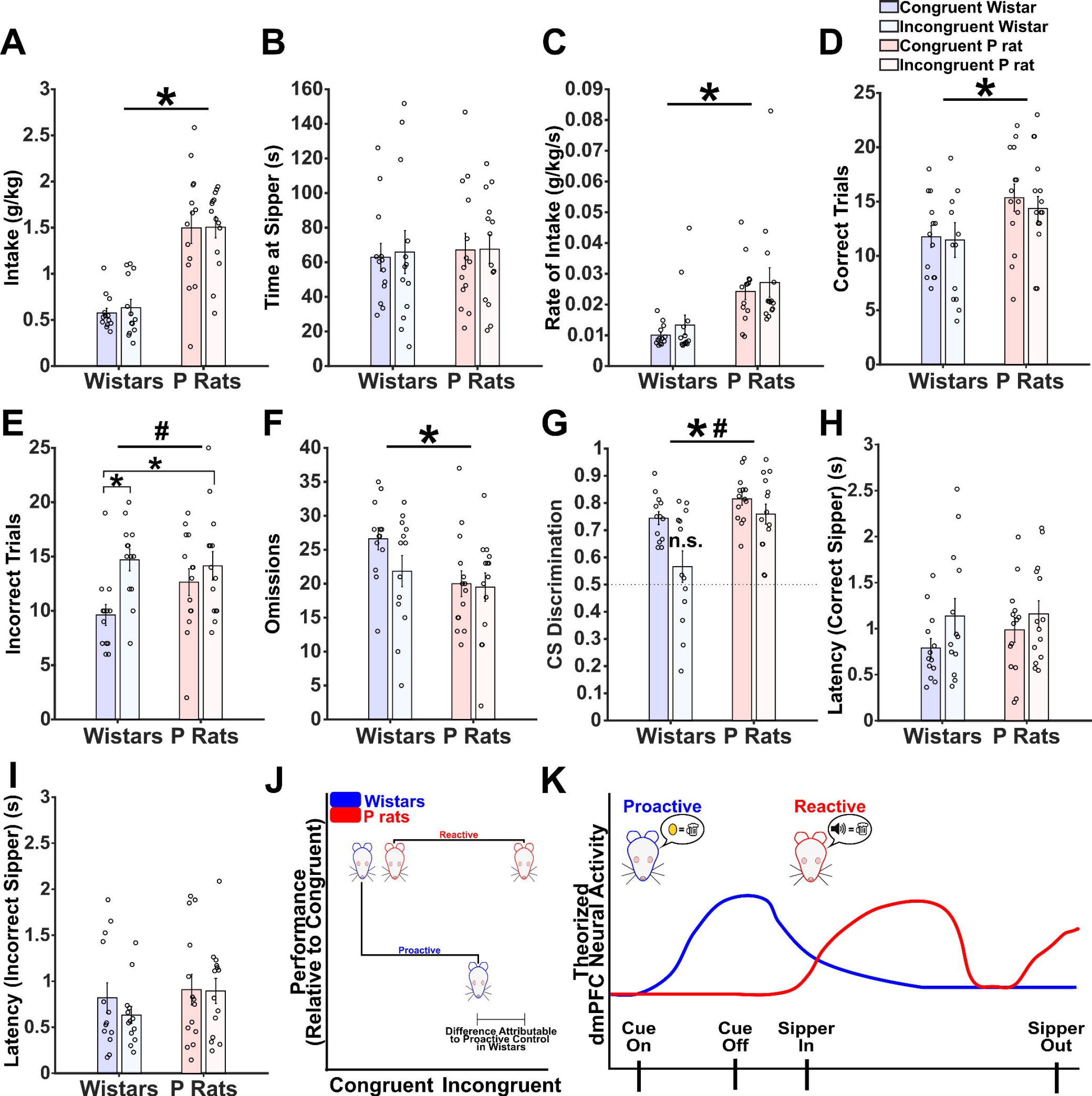
Incongruent sessions disrupt Wistar performance. A. Intake in g/kg is shown for each strain and session type. A main effect of strain (**\***) was detected. **B.** Sipper occupancy was quantified and no main effects of either strain or session type was detected. **C.** Rate of intake is depicted and a main effect of strain (*) was detected. **D.** Correct trials were determined per session and a main effect of strain (*) was detected. **E.** Incorrect trials were determined per session and a main effect of session type (#) was detected. Post hoc tests indicated that congruent Wistars made fewer mistakes than incongruent Wistars and incongruent P rats. **F.** Omissions were determined per session and a main effect of strain (*) was detected. **G.** CS Discrimination was calculated and a main effect of strain (*) and session type (#) were detected. Additionally, incongruent Wistars’ ratio was not different from 0.5. **H.** Latency to approach the correct sipper is shown. A trending main effect of session type was found. **I.** Latency to approach the incorrect sipper is shown. No main effects of strain or session type was found. A MANOVA detected significant differences between congruent and incongruent sessions in Wistars but not P rats. This suggests an overall difference in behavioral performance exists in Wistars but not P rats when the session contingency is changed. **J.** Mockup model data that reflects the results of the MANOVA. **K.** Theoretical model of our hypothesis that neural activity in Wistars will precede alcohol delivery whereas neural activity in P rats will coincide with alcohol delivery.

While measures of alcohol consumption did not differ between session types, Wistars showed behavioral differences related to the incongruent sessions. Correct approaches are defined as approaching where the sipper has descended on the first attempt. These data show that there is a main effect of strain (ANOVA, F(1,50) = 6.572, p = 0.013, **Figure 2D**) on the number of correct approaches. Interestingly, Wistars in the incongruent sessions made an increased number of incorrect approaches. A main effect of session type was detected (F(1,50) = 8.129, p = 0.006, **Figure 2E**). Post-hoc analyses indicated that congruent Wistars made significantly fewer mistakes compared with both incongruent Wistars (Tukey’s HSD: p = 0.018, 95% CI: [-5.077: -0.662]) and incongruent P rats (Tukey’s HSD: p = 0.0375, 95% CI: [-4.528, - 0.193]). Wistars also made more omissions (ANOVA, F(1,50) = 5.042, p = 0.029, **Figure 2F**). In sum, the data indicate that incongruent trials have a modest, impairing effect on Wistars’ performance that is not detectable in P rats.

If the animal detected a change in the CS-reward contingency, this might also indicate that meaning of the CS+/- was altered. The ratio was the amount of CS+ approaches divided by the sum of CS+ and CS- approaches. CS+ approaches included both correct and incorrect approaches. Values near 0.5 indicate that the animal approaches both CS types at equal rates. A main effect of both strain (ANOVA, F(1,50) = 12.280, p < 0.001) and session type (F(1,50) = 9.613, p = 0.003, **Figure 2G**) were observed. A single sample t-test indicated that each distribution was significantly different from 0.5 except Wistars during incongruent sessions (t (12) = 1.141, p = 0.276) suggesting that the incongruent contingency uniquely interfered with Wistars’ CS discrimination.

We additionally analyzed latency to the sipper. The latency measure begins as soon as the sipper descends. Correct latencies required the animals to visit the correct sipper first. A trend of session type was detected (F(1,50) = 3.18, p = 0.081, **Figure 2H**) where incongruent sessions may lead to longer latencies. No main effects were observed in the latency to reach the incorrect sipper (**Figure 2I**).

To determine the extent to which Wistars or P rats display global impairments due to incongruent sessions, a MANOVA was conducted for each strain. All variables from **Figure 2** were included except for the omissions due to constraints on collinearity. Overall, Wistars’ behavior was significantly altered by incongruent sessions (MANOVA, F(1,24) = 3.452, p = 0.015) whereas P rats do not show global alterations due to the incongruent sessions (MANOVA, F(1,26) = 0.557, p = 0.800). This suggests that Wistars are impacted more by the contingency change than P rats as evidenced by the session-wide metrics. **Figure 2J** depicts a mockup, hypothetical figure based on the results of the MANOVA. In general, we see deficits during the incongruent session in only Wistars. This led to the hypothesis that they are utilizing different cognitive control strategies from P rats. **Figure 2K** is a theoretical figure that graphically depicts our hypothesis that neural activity in Wistars will precede alcohol delivery whereas neural activity in P rats will coincide with alcohol delivery. This is due to the timing of goal representations in each strain; Wistars employ proactive cognitive control that maintains the goal over time whereas P rats employ reactive cognitive control that generates a representation of the goal only when alcohol is available. To find further evidence of our hypotheses, the temporal aspects of Wistars’ mistakes, both across and within trials, was analyzed next.

### Wistars make immediate mistakes during incongruent sessions

Wistars show deficits in the aggregate session data. We next sought to determine when within a session they show these deficits. The mean probability to approach the correct or incorrect sipper was determined for each strain and session type. Correct approaches only include those where the animal successfully approaches the alcohol-containing sipper on the first attempt. The mean probability was calculated as a 3-trial moving average. In Wistars, a main effect of session type (RANOVA, F(1,48) = 4.097, p = 0.0485) and an interaction between the session type and trial accuracy was observed (F(1,48) = 5.373, p = 0.025, **Figure 3A**). On early trials Wistars exhibit decreases in correct approaches that are paralleled by an increase in incorrect approaches. This suggests that they are implementing their previously learned rule. In P rats, no between or within subject main effects or interactions were found. Critically, no difference in session type was observed for P rats (RANOVA, F(1,52) = 0.056, p = 0.814, **Figure 3B**). Ultimately the data suggests meaningful differences in the session-wide time course of behavior in Wistars emerges as a function of the change in contingency between the CS+ and reinforcer.

**Figure 3.**
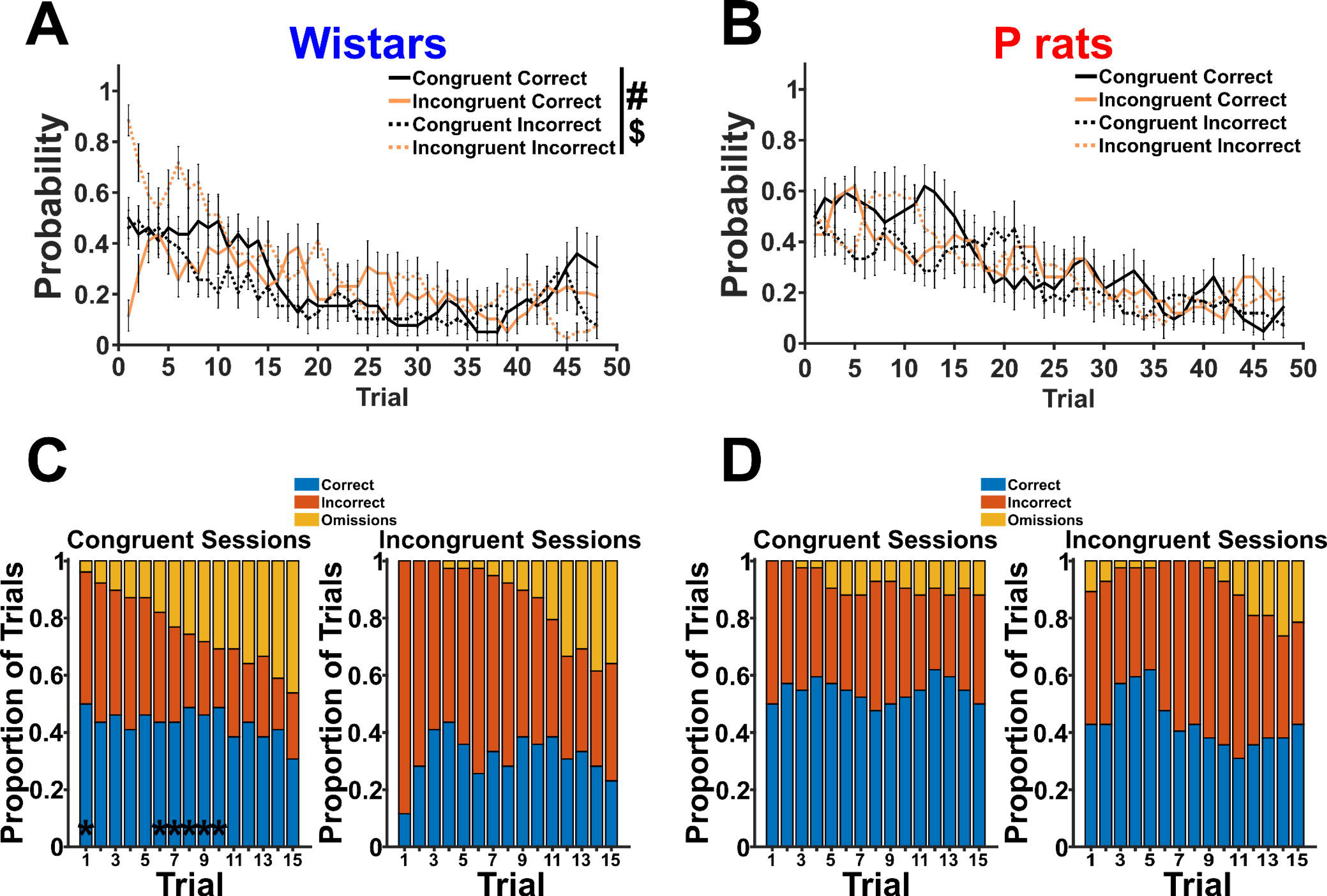
Wistars make mistakes earlier within a *session*. A. The probability to make a correct or incorrect approach is shown for each session type for Wistars. A main effect of trial type ($) and a main effect of session type (#) was detected. Critically, Wistars make more mistakes earlier in the session. **B.** The same information from **A** is shown but for P rats. Critically, neither a main effect of trial type or session type was detected. **C.** The mean proportions of correct, incorrect, or omissions is shown for the first 15 trials in Wistars. Asterisks (*) denote trials where the proportions from the incongruent sessions are significantly different than the proportions from the congruent sessions. **D.** The same information as **C** is shown but for P rats.

Next, the difference between mean proportions of correct, incorrect, or omissions in the first 15 trials for was analyzed for each strain using a chi-squared test. Asterisks on the congruent session graph denote trials where the proportions between congruent and incongruent sessions significantly differ after FDR-corrections. Ultimately, we see that Wistars (**Figure 3C**), but not P rats (**Figure 3D**), are uniquely affected by the changed contingency early within a session.

After finding clear evidence that Wistars are affected early within a session, we next sought to determine when within a trial they were affected. We found that Wistars are impacted in the first few trials, where they reach the correct sipper less, and reach the incorrect sipper immediately after its entry (**Figure 4A**). Conversely, P rats are diffusely affected throughout the first 15-trials (**Figure 4B**). Next, we quantified the differences between sipper occupations. First, we divided time into 5 epochs (Pre-Cue, Cue On, Early Alcohol Access (E1), Late Alcohol Access (E2), and Post-Access). Next, the maximum value per epoch per trial was found and averaged resulting in the maximum change in likelihood per epoch. At the correct sipper, no main effects or interactions were detected (**Figure 4C**). Next, the maximum difference in probability that each respective strain was at the incorrect sipper during that epoch was found. A significant main effect of strain (F(1,28) = 4.792, p = 0.037) and a significant interaction between strain and epoch (F(1,28) = 6.238, p = 0.019) was detected (**Figure 4D**). Post-hoc analyses indicated that a significant difference between Wistar and P rats was detected in the E2 epoch (p = 0.007, 95% CI: [0.017, 0.097]). While early access and cue-onset periods were not different according to the post-hoc test, restricting the analyses to the first 5 trials would likely show that the Wistars are more likely to error early within a trial compared with P rats.

**Figure 4.**
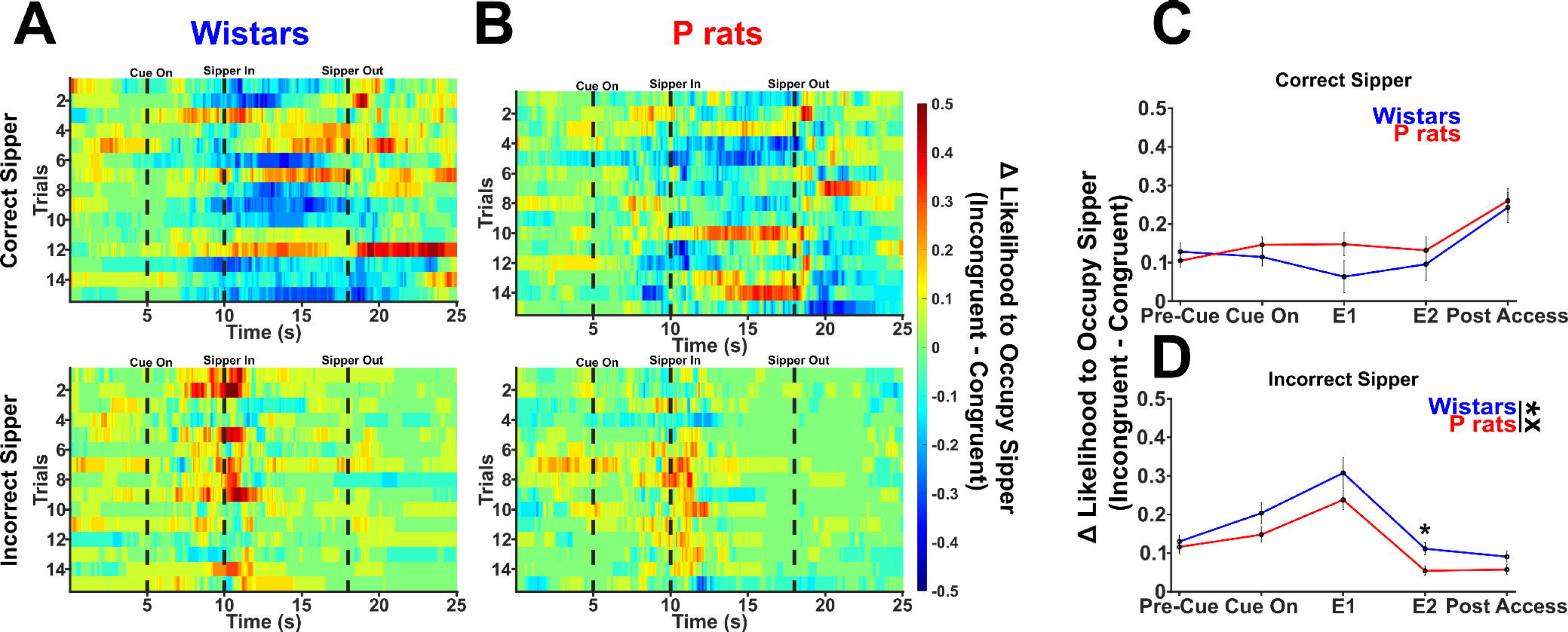
Wistars make mistakes earlier within a *trial*. A. The change in sipper occupancy probability is depicted for both the correct and incorrect sipper in Wistars. Warmer colors indicate a higher probability that the sipper was occupied during incongruent sessions. Cooler colors indicate a higher probability that the sipper was occupied during congruent sessions. Green indicates no differences between congruent and incongruent. Critically, Wistars are more likely to visit the incorrect sipper during incongruent sessions early within the trial and early within the session. However, as the session progresses, the difference between P rats and Wistars disappears. **B.** The same information as **A** is depicted except for P rats. P rats are not as likely to immediately visit the incorrect sipper during incongruent sessions. However, their performance degrades as the session goes on. **C.** The change in probability is quantified for the correct sipper. Trials were broken into 1 of 5 epochs: Pre-Cue [-4s - 0s], Cue On [0s – 5s], E1 [5s – 9s], E2 [9s – 13s], and Post Access [14s – 18s]. E1 and E2 correspond to early and late alcohol access. The maximum value per trial was obtained for each epoch and then averaged across trials. **D.** The change in probability is quantified for the incorrect sipper. The asterisk beside the figure legend indicates the main effect of strain. The x indicates the significant interaction between strain and epoch. The asterisk above E2 indicates the post-hoc difference at that epoch between Wistars and P rats.

The behavioral data suggests that differences emerge between P rats and Wistars at both a baseline level and in how they solve the task when the contingency has been altered. Wistar rats appear to utilize a task schema that guides their behavior proactively according to the learned set of rules the task affords. When the contingency of the task changes in the incongruent sessions, they then have difficulty adapting presumedly because they are using the previously learned rule. P rats, however, solve the task through other means. These behavioral data support our overarching hypothesis that Wistars utilize proactive cognitive control whereas P rats utilize reactive cognitive control.

### Wistars show differences in cue encoding while P rats show differences in alcohol access encoding

**Figure 5** describes the data structure prior to PCA. No differences were found in the raw, averaged firing rates (**Figure 6A**). However, a weak signal surrounding the cue onset for Wistars and strong signals surrounding the sipper descent and ascent for P rats can be qualitatively observed. PC projections of the top 5 variance explaining PCs for Wistars show that signals that were obscured in the mean become more prominent in the principal component projections. For Wistars during congruent sessions, the PC projections show large amounts of cue-locked activity (**Figure 6B**). However, the contingency change selectively blunts this cue encoding in Wistars only (**Figure 6D**). P rats in the congruent session intriguingly show no signals around the onset of the cue and instead the top 5 PCs are focused on the sipper descent and ascent which signals the start and end of alcohol availability (**Figure 6C**). In P rats, this alcohol-onset related activity is exaggerated during incongruent sessions (**Figure 6E**).

**Figure 5.**
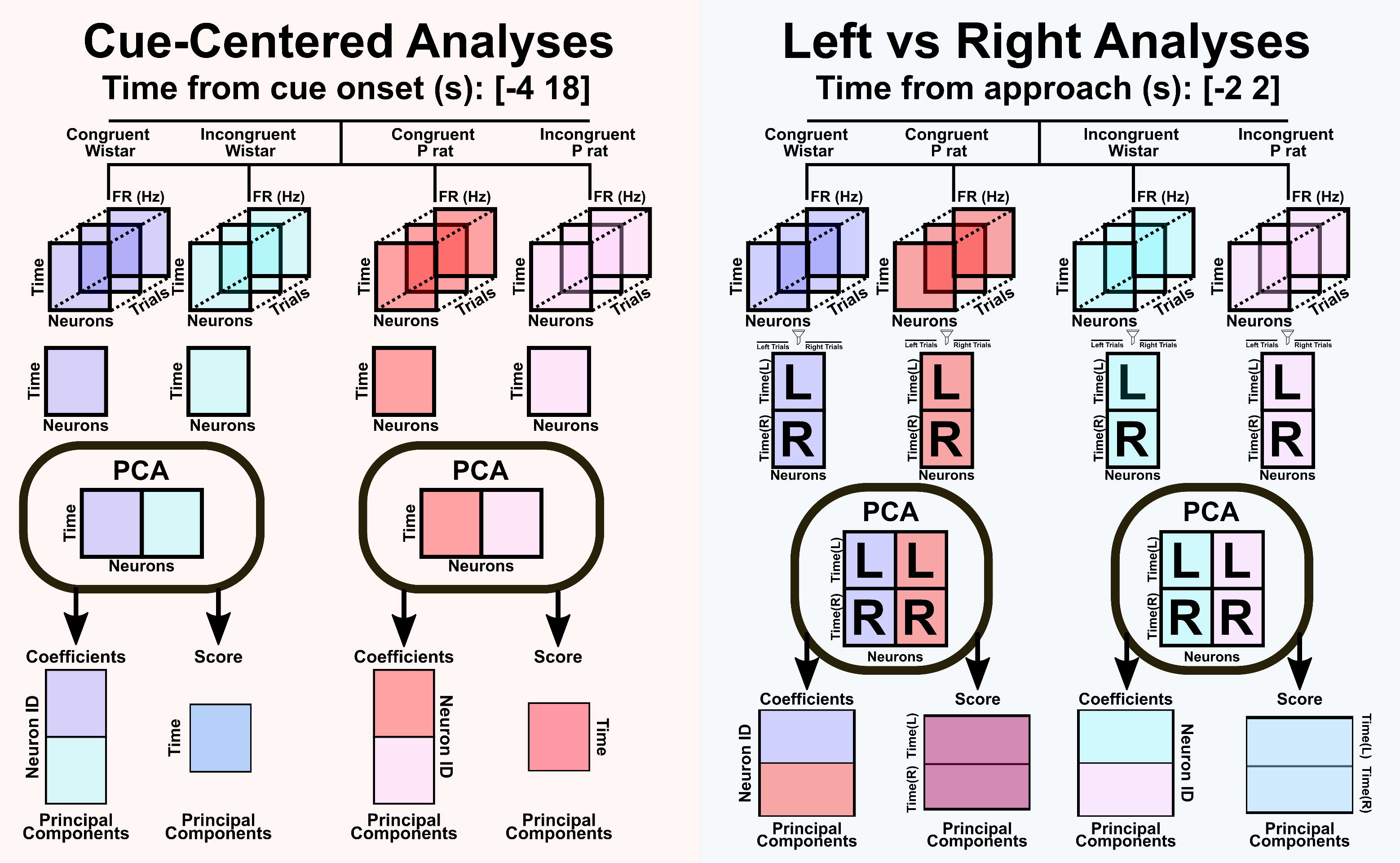
PCA methodologies for cue-centered and approach-centered analyses. Cue-centered PCA analyses were accomplished by first smoothing 100ms binned firing rates with a gaussian kernel. Then a NxMxT matrix was created where N refers to the number of neurons, M refers to the number of timepoints, and T refers to the number of trials. 4 seconds prior to the cue onset and 18 seconds after the cue onset were analyzed [-4 18] resulting in 22 total seconds. At 100ms bins, this resulted in 221 time bins. The number of trials, T, were the first 15 trials of the session. The mean of trials was calculated such that a NxM matrix remained. The NxM matrix for the congruent and incongruent Wistar sessions were vertically concatenated separately from the P rat sessions. PCA was then performed and the resulting projections and coefficients were analyzed. In the approach-centered analyses, everything was the same except that the number of time bins was greatly reduced and centered on the approach rather than the cue onset. In the left versus right analyses, up to 15 correct trials were averaged. All correct left trial means were averaged separately from all correct right trial means. Then the left and right trial means were horizontally concatenated which doubled our M dimension. Congruent Wistar and congruent P rat sessions were then vertically concatenated. The same was done with incongruent Wistar and incongruent P rat sessions. We then ran PCA to obtain the coefficient and score products. Following this, all PCA products were filtered into left versus right.

**Figure 6.**
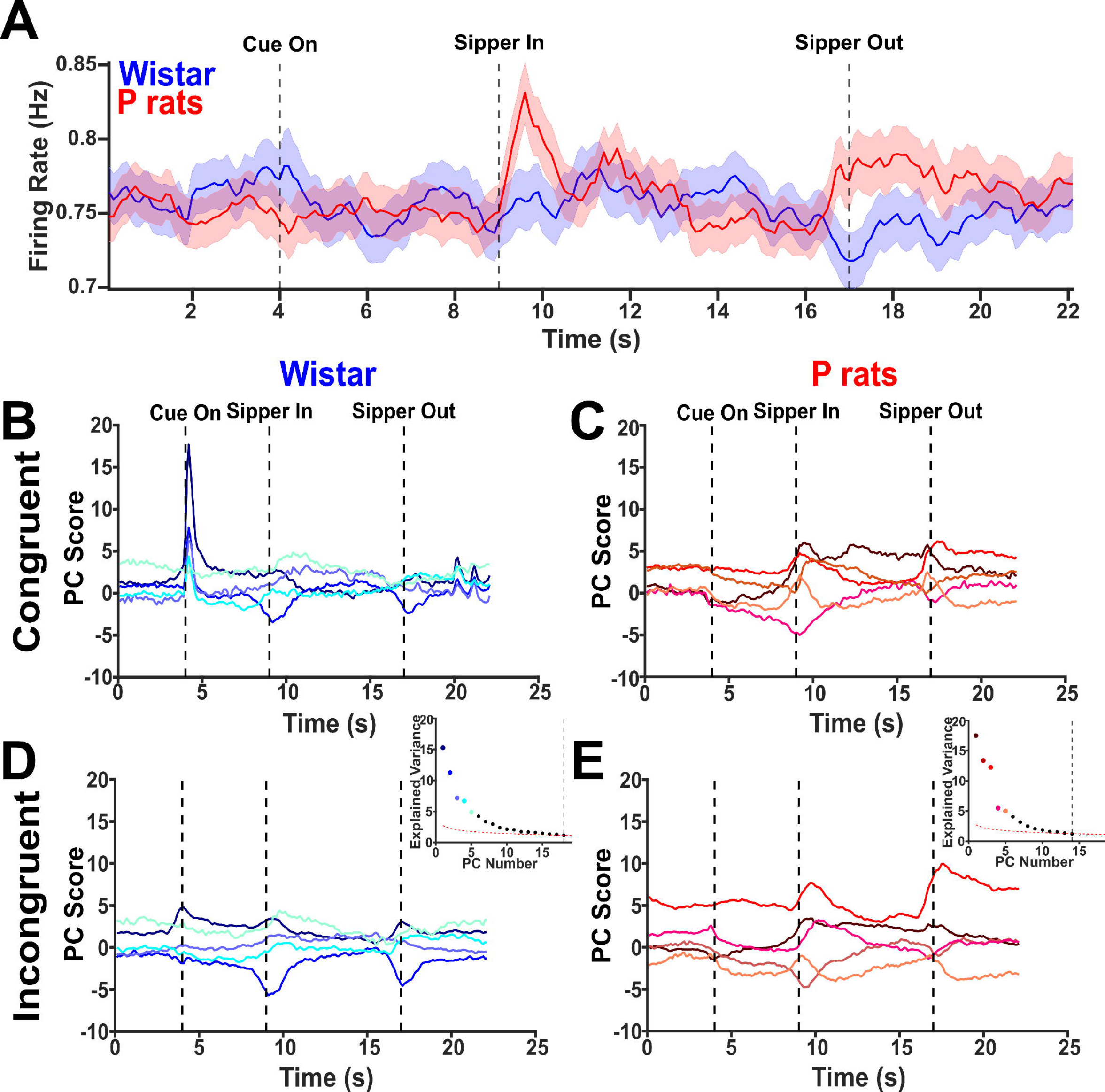
Principal components highlight several differing features between strains and session types. A. The mean firing rates for each strain collapsed across session type is depicted. **B.** The top 5 PCs for Wistars performing the congruent sessions is shown. Additionally, the inset describes the explained variance of each, the dashed line indicates the broken stick point. Interestingly, several of the PCs encode the onset of the cue. **C.** The same information as **B** except for P rats is shown. Several of the PCs encode the sipper descent and ascent. **D.** The top 5 PCs for Wistars performing the incongruent sessions is shown. Cue encoding during incongruent sessions is blunted in Wistars. **F.** The same as **D** but for P rats. PC3 shows enhanced sipper encoding during the descent and ascent.

Overall, the first 5 PCs demonstrate an overview of the neural activity in these tasks, and it indicates that neural activity in Wistars is more robustly modulated by the CS and is blunted during incongruent sessions. Conversely, P rat neural activity is robustly modulated during the periods of time corresponding to alcohol access: the sipper in and sipper out, and this is moderately altered during the incongruent session.

### Wistars show left/right encoding prior to approaches, P rats show left/right encoding after approaches

Next, we hypothesized that Wistars would show greater spatial encoding prior to their approaches compared with P rats. Differences in spatial encoding suggest that each side, left or right, has a corresponding strategy representation reflective of prospection. Greater distances in a PC space would then suggest that each side has a well-developed internal rule guiding subsequent behavior. Dedicating more cognitive resources to differentiate these two sides suggests a greater reliance on proactive control.

To determine the encoding of left versus right strategies, the data were organized differently from the first analysis prior to the PCA. First, only correct approaches were utilized where the animal successfully approached the alcohol-containing sipper on the first attempt. Next, we found all correct approaches for the left sipper and all correct approaches for the right sipper. Then we averaged all left approaches separate from all right approaches. In general, the raw data show no major differences across conditions (**Figure 7A,B**). Left and right choices were then concatenated in time such that a Nx2M matrix of the trial averaged firing rates was created where N was each neuron and M was each timepoint. The data were then split by session type such that all congruent sessions were run together, and all incongruent sessions were run together, regardless of strain. PCA was performed on this Nx2M matrix and the PCA products were indexed such that left and right choices were separated once more (see **Figure 5** for overview). **Figure7C, D** shows the PC projections of the top 5 PCs with solid lines indicating left correct choices and dashed lines indicating right correct choices. For congruent sessions, we included the top 9 PCs and for incongruent sessions we included the top 8 PCs. We first sought to determine if there were any gross changes in the distance between left and right choices. **Figure 7E** describes the data analysis process. In brief, we permuted over every possible 3-dimensional PC space for each session type. For congruent sessions, this resulted in 84 possible permutations and for incongruent sessions this resulted in 56 possible permutations. Within each permutation, we randomly selected 1000 individual left and 1000 individual right trials from the possible pool of each and calculated the PC space distance between each of these trial combinations prior to the approach time. We then took the mean of those distance combinations to determine the mean distance of each permutation. An overall group difference was observed in the distances of each conditions left/right encoding (Kruskall- Wallis (3, 276), χ^2^ = 67.22, p < 0.001, **Figure 7F**). Tukey-Kramer multiple comparisons were used to follow up on the overall group differences. The distance between left and right trials in the Wistar, congruent sessions were significantly greater than the Wistar, incongruent sessions (95% CI: [69.619, 141.393], p < 0.001), the P rat, congruent sessions (95% CI: [40.306, 104.503], p < 0.001) and the P rat, incongruent sessions (95% CI: [38.154, 109.929], p < 0.001). No other significant differences between groups were found, including between P rat congruent and incongruent conditions. Spatial encoding is therefore higher in the Wistars in congruent sessions, and, subsequently it becomes less accurate during incongruent sessions, as evidenced by both their performance and neural encoding.

**Figure 7.**
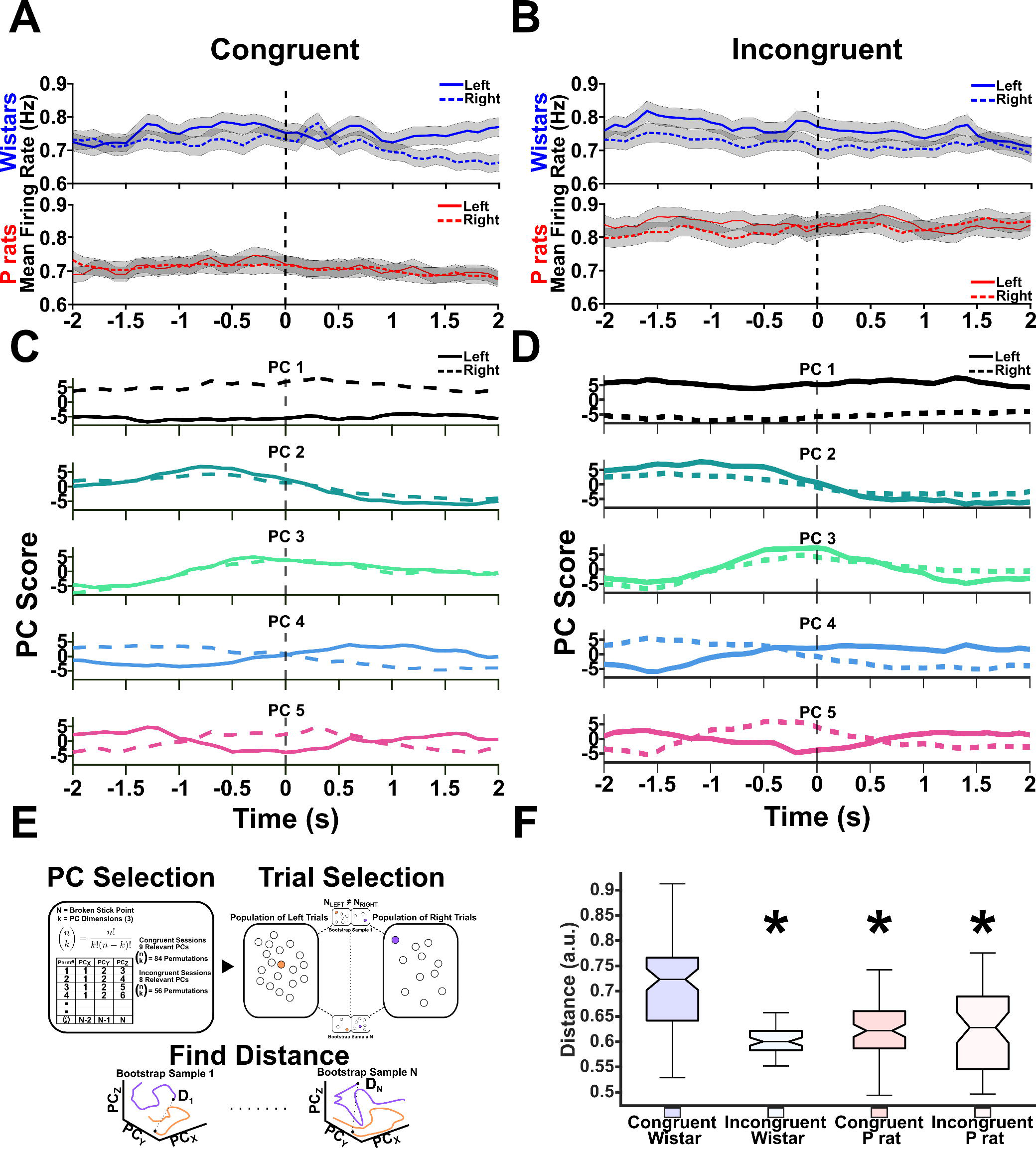
Wistars show greater spatial encoding. A & B. The mean firing rates for left and right choices are depicted. **C & D.** The top 5 PCs for congruent and incongruent sessions are shown. Solid lines depict left trials, dashed lines depict right trials. **E.** A description of the analysis performed to determine the encoding differences between left and right trials. In brief, several permutations in a 3-dimensional PC space were identified. In each permutation, 1000 random left and right trials were compared. The mean of those 1000 trials was then take for each permutation. **F.** The results of the permutation analysis are shown. Briefly, the mean of the bootstrap samples was found for each permutation. The mean was then taken prior to approach to determine separation of left versus right choices prior to approach initiation. An overall effect was detected. * Tukey-Kramer post-hoc tests identified that the distance between left and right trials in the Wistar congruent session was higher than every other strain and session type condition.

To interrogate when within a trial P rats and Wistars differentially encode left versus right choices, we employed an additional analysis that allowed us to look at the raw firing rates of each population. In brief, neurons were indexed based on whether they were positive or negative loaders on each PC. Once again, we used a threshold of greater than/less than or equal to 0.01/-0.01. From this index, we pooled together neurons with the same signs from each condition and determined their means. This resulted in time varying mean firing rates for both positive and negative loading neurons in each direction (left vs. right) as well as each session type. To focus our analysis, we sought to identify PCs that selectively encoded left and right choices and varied over the approach period. A RANOVA was conducted on the mean firing rates that were positively or negatively loaded onto each PC. PC4 solely demonstrated a significant time by side interaction in both the congruent and incongruent sessions. Qualitatively, PC4 exhibited similar characteristics in both session types.

Because of the PCs unique characteristics, we chose to further investigate it (**Figure 8A, B** for Wistars in congruent and incongruent sessions, respectively, **Figure 8C, D** for P rats in congruent and incongruent sessions, respectively). We were then interested in the firing rate difference between left and right choices before and after the approach. Large firing rate differences between left and right choices are then interpreted as an index of selective, spatial encoding. We were able to determine this by taking the difference in firing rate between each side and averaging the first half prior to approach and then averaging the second half after the approach. Positive values indicate that the mean of the condition is skewed towards encoding the left whereas negative values indicate that the mean of the condition is skewed towards the right. A two factor, repeated measures ANOVA was used where approach coded for whether the animal was before or after their approach. A significant main effect of approach epoch was found (F(1,76) = 519.468, p < 0.001) and a significant main effect of session type was found (F(1,76) = 10.849, p = 0.002, **Figure 8C**). For negative loading neurons, an additional main effect of approach was found (F(1,76) = 642.096, p < 0.001). Interestingly, a main effect of strain (F(1,76) = 43.170, p < 0.001) but not session type was found. A significant interaction between strain and session type was also found (F(1,76) = 4.021, p = 0.049, **Figure 8D**).

**Figure 8.**
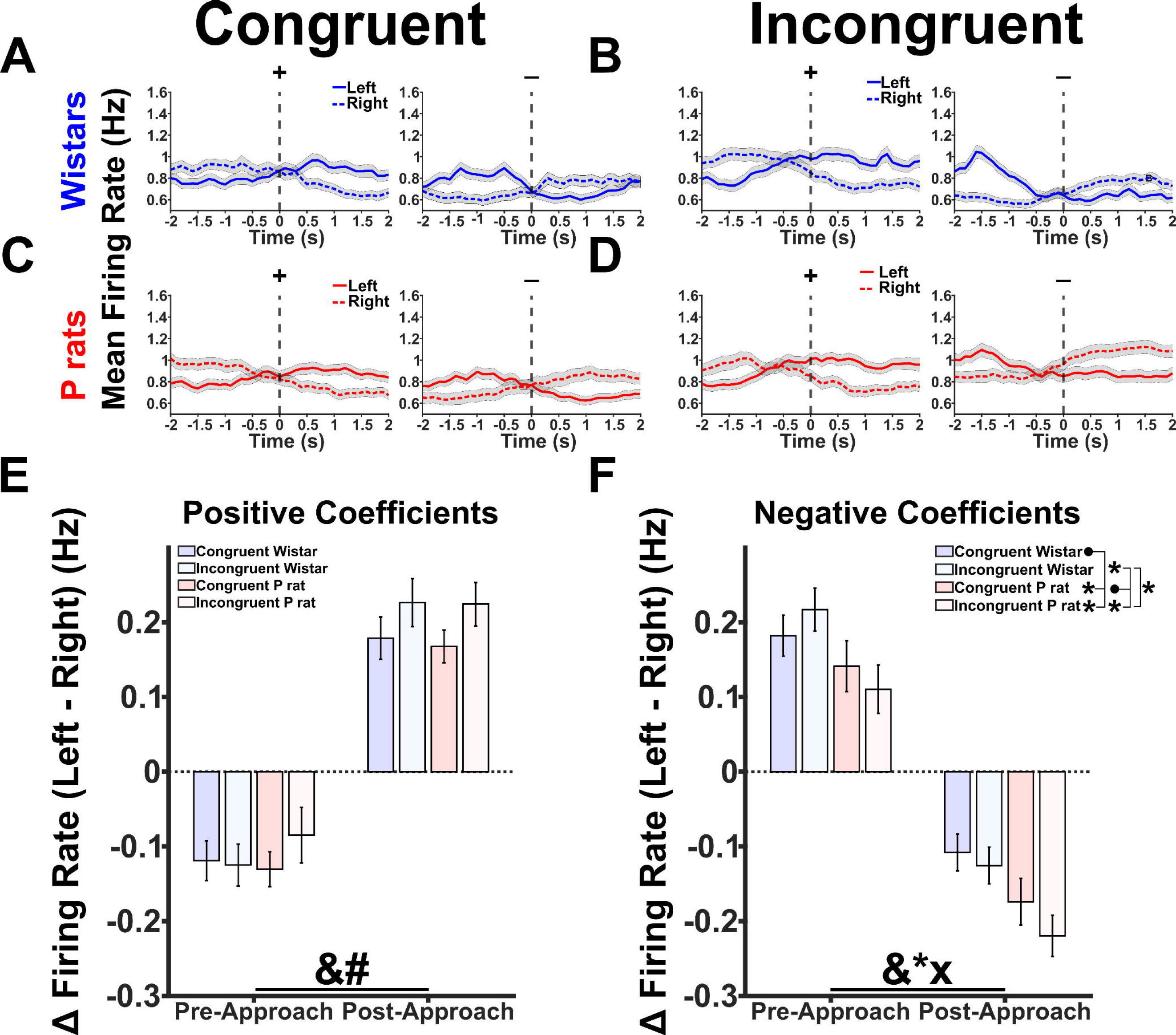
Wistars and P rats show differential spatial encoding pre-and-post approach. **A & B** show the mean firing rate for neurons that either positively or negatively load onto PC4 in both the Congruent and Incongruent sessions for Wistars. Critically, the mean firing rate diverges at or prior to the time when each animal approaches making it a useful index of neural activity. **C & D** show the same mean firing rate for positive and negative loading neurons for PC4 in the P rat neural population. The same divergent feature at the approach time emerges. **E.** The difference between left and right firing rates was calculated and compared across each session type and strain combination before and after the approach for positive coefficients. A main effect of approach (&) and session type (#) was found. **F.** The same analysis as **E** but for the negative coefficients. A main effect of approach (&) and strain (*) was found. Additionally an interaction between strain and session type was found. Further probing this effect showed that the congruent Wistar groups were significantly different from the congruent P rat and incongruent P rat groups. The congruent P rat group were significantly different from the incongruent Wistar and incongruent P rat group, and the incongruent Wistar group was significantly different from the incongruent P rat group.

Probing this interaction shows that congruent Wistar sessions differed from congruent P rat sessions (t(76) = 3.228, p = 0.006), incongruent P rat sessions (t(76) = 5.541, p < 0.001), but not incongruent Wistar sessions (t(76) = -0.523, p = 0.603). Congruent P rat sessions differed from incongruent P rat sessions (t(76) = 2.313, p = 0.047), and incongruent Wistar sessions (t(76) = -3.751, p = 0.001). Last, incongruent P rat and incongruent Wistar sessions differed from one another (t(76) = -6.064, p < 0.001). Taken together, the results of **Figure 8** suggest that two competing populations of neurons exist that differentially encode left versus right choices in both P rats and Wistars. The positive loading population is selective for session type whereas the negative loading population demonstrates approach sensitivity; specifically, Wistars show greater differences in left versus right firing rates prior to the approach whereas P rats show greater differences after the approach. This result is in line with our previous PC space analysis and further supports our hypothesis of Wistars exhibiting proactive cognitive control whereas P rats exhibit reactive cognitive control.

### Left versus right encoding distances are correlated with markers of task performance

Last, we sought to determine if the distances between left and right trial trajectories had meaningful correlations with behavioral metrics found in **Figure 3**. We hypothesized that stronger representations of side information should lead to faster reaction times on congruent trials. However, if side encoding is driving behavior this relationship should be flipped on incongruent trials with higher distances corresponding to longer reaction times. We fit a linear model to both the Wistar data and P rat data to further test if distance significantly predicted latency. For Wistars, the fit equation was: Latency = 0.924 – 0.338 * (Distance) – 0.427 * (Session) + 1.839 (Distance * Session). The overall regression was statistically significant (R^2^ = 0.408, F(1,21) = 4.83, p = 0.010). Neither distance (β = -0.338, p = 0.327) or session (β = - 0.427, p = 0.152) significantly predicted latencies. However, the interaction between distance and session was a significant predictor of latencies (β = 1.839, p = 0.004). A non-significant negative correlation was observed for congruent sessions (Pearson’s r = -0.363, p = 0.223) while during incongruent sessions, Wistars had a significant, positive correlation (r = 0.657, p = 0.020, **Figure 9A**). In our linear model for P rats the fit equation was: Latency = 0.950 + 0.108 * (Distance) – 0.102 * (Session) + 0.822 * (Distance * Session). The overall model was not statistically significant (R^2^ = 0.223, F(1,22) = 2.11, p = 0.128). Distance (β = 0.108, p = 0.759),

**Figure 9.**
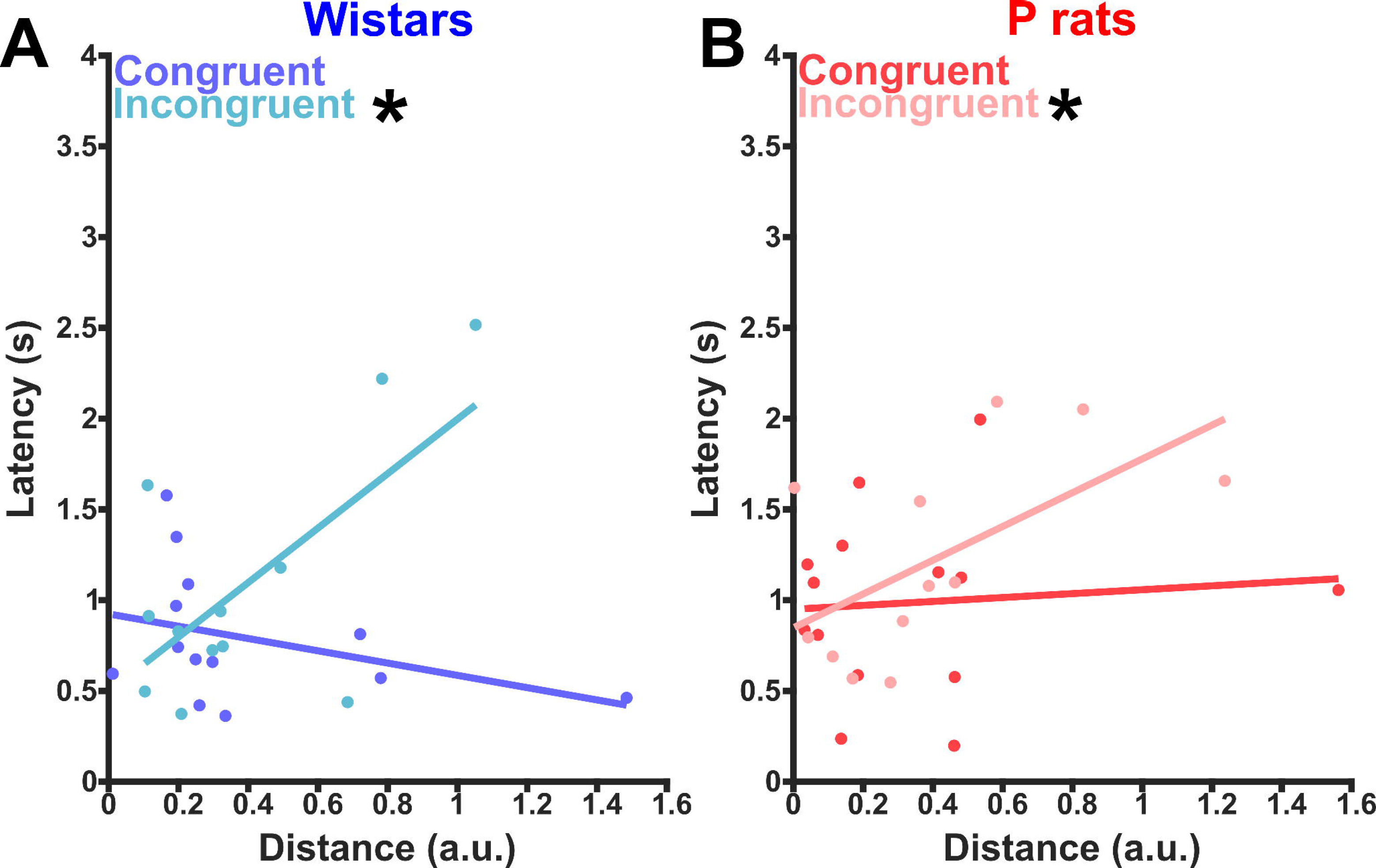
Spatial encoding is meaningfully related to latency to approach the sipper. A. The correlation between latency to approach the sipper and the distance between left and right trials for each session is shown. In the congruent condition, latency and distance are non-significantly and negatively correlated whereas in the incongruent conditions latency and distance are significantly positively correlated. Overall, the Wistars show a significant interaction between distance encoding and session type. This suggests that it is beneficial to encode side prior to moving during congruent sessions but may be detrimental during incongruent sessions. **B.** The same information as was seen in **A** but for P rats. In the P rats, in the congruent condition latency and distance are non-significantly and positively correlated whereas in the incongruent condition latency and distance are significantly and positively correlated.

Session (β = -0.102, p = 0.723) nor the interaction (β = 0.822, p = 0.147) significantly predicted latency. Interestingly, P rats did have a slight, significant positive correlation during incongruent sessions (Pearson’s r = 0.590, p = 0.043, **Figure 9B**). Taken together, these results suggest that markers of task performance are linked with our distance metric. In congruent sessions this suggests it *may* be beneficial to robustly encode the task set with greater distance values resulting in lower latencies. On the contrary, the sign change in the incongruent sessions suggests that it is detrimental to robustly encode the task set.

## Discussion

The present data provide evidence that P rats and Wistars utilize different strategies when seeking alcohol in the 2CAP task. It was hypothesized that Wistars’ behavior is best described as being dominated by proactive cognitive control whereas P rats’ behavior is best described as being dominated by reactive cognitive control. Behavioral and neural evidence indicates that Wistars are prospectively engaged in the task while P rats tend to engage reactive control in response to environmental stimuli. This difference in cognitive control styles may be critical for why the two strains diverge in their drinking behaviors.

2CAP is a model of alcohol seeking in response to cues. Several studies utilizing it have been published by our lab (McCane et al., 2014; Linsenbardt & Lapish, 2015; Linsenbardt et al., 2019; Timme et al., 2020; Timme et al., 2022) and demonstrate strain differences between Wistar and P rats. Several of these studies have also examined changes in the alcohol contingency whether it be through the addition of quinine (Timme et al., 2020; Timme et al., 2022) or the substitution of alcohol with water (Linsenbardt & Lapish, 2015; Linsenbardt et al., 2019). In both cases, P rats continue to demonstrate strong seeking behaviors despite changes in the contingency. However, a novel angle of the present study is measuring neural activity while disrupting the contingency that predicts reward. In the aggregate, we see clear signals that Wistars are differentially affected by the changed contingencies that are not as apparent in P rats.

In the present dataset and in previous work (Timme et al., 2022) we observed that the CS+ elicits approach behavior in both strains whereas the CS- does not. This is evidence that the animals have formed an association between the presentation of the CS+ and the subsequent availability of alcohol. However, as evidenced by the present data, the specific sequence of actions the CS+ elicits likely differ between and within a strain. For at least the majority of Wistars, they can follow the cue light to the detriment of task performance in the incongruent sessions. Whereas for P rats, the meaning of the CS+ seems less certain. An alternative explanation for the purpose of the CS+, particularly for animals exhibiting reactive cognitive control, may be as an occasion setter that guides attention to the more proximal cue of the noise from the sipper motors (Fraser & Holland, 2019). However, because our sipper motors are active on both sides of the 2CAP chamber, the noise is not a discriminative stimulus which may explain why the P rats’ performance is consistently at chance.

Other rodent studies have explored proactive and reactive cognitive control. Most of these have used tasks that measure perceptual decision making and stress psychophysical measures such as reaction times. Longer reaction times often reflect reactive control processes as the agent gathers evidence (Gu et al., 2020; Hernández -Navarro et al., 2021). Shorter reaction times indicate a task set exists that is stimulus independent (Hernández -Navarro et al., 2021). Additionally, several parallel brain regions are involved in proactive and reactive computations such as the globus pallidus (Gu et al., 2020) and red nucleus (Brockett et al., 2020). Together, these data suggest that proactive and reactive cognitive control processes are widely distributed and competing processes. Network approaches would likely reveal differences in proactive and reactive cognitive control between P rats and Wistars.

While the current study did change the behavioral task in a way that can be considered a true “reversal”, there are points of contact with reversal tasks. Bartolo & Averbeck (2020) found that prefrontal cortex activity predicts state switches in monkeys at rates faster than what would be expected in typical reinforcement learning scenarios. Furthermore, Dalton and colleagues (2016) suggest that rodent dmPFC regions are critical for monitoring action-outcome contingencies and Atilgan and colleagues (2022) found evidence that mice in a probabilistic reversal learning task learn the structure of the task and that dmPFC activity is related to abrupt changes in task performance, similar to work performed by Bartolo & Averbeck (2020) as well as Durstewitz and colleagues (2010). Ultimately, the data suggest that the dmPFC is critical for generating task sets. Deviations from the expectancies are tracked in the dmPFC in a proactive manner that allows for flexible and fast switching behaviors. However, specific cognitive demands are required for the dmPFC to be critical for reversals (Hamilton & Brigman, 2015).

Understanding how model-free or model-based learning systems interact with proactive and reactive cognitive control is a critical future direction.

Cognitive control is thought to be linked with the dopamine (DA) system (Carter & Cohen, 1998; Schultz, 1998). DA has a critical role in the perception of incoming information and how those percepts, such as cues, lead to actions (Cools & d’Esposito, 2011; Jacob et al., 2013). A clinical delineation in cognitive control task performance in those with Parksinson’s disease versus those with Huntington’s disease (HD) suggests that prefrontal DA is critical for cognitive control (Unschuld et al., 2012; Timmer et al., 2018; Júlio et al., 2019). Differences in cortical DA may explain why P rats and Wistars differ in cognitive control. A possible mechanism for the difference between strains is the differential expression of the enzyme catechol-O-methyl transferase (COMT) gene (McCane et al., 2017). COMT has an active role in the clearance of dopamine from the dmPFC. COMT was observed to be under expressed in the dmPFC of male P rats (McCane et al., 2017). While P rats generally have less extracellular DA (Engleman et al., 2006), 2CAP’s cues may evoke and maintain greater cortical DA given the lack of COMT clearance thereby invigorating reward responding and motivation. Additionally, the COMT inhibitor, Tolcapone, was shown to suppress alcohol intake in P rats (McCane et al., 2014). Together, this suggests that differences in the presence and clearance of dopamine in the dmPFC is critical for alcohol seeking phenotypes. An additional mechanism for the differences observed may be the lack of metabotropic glutamate receptor 2 (mGluR2) in P rats (Zhou et al., 2013). mGluR2 is a diffuse and inhibitory receptor (Ohishi, et al., 1993; Schoepp et al., 1992). Long-term exposure to alcohol has been shown to cause decreases in mGluR2 expression in Wistars and humans (Meinhardt, 2013) that can be rescued through viral upregulation (Meinhardt, 2013). In Wistars and Long-Evans, selective activation of mGluR2 can also prevent cue-induced relapse to alcohol seeking (Bäckström & Hyytiä, 2005; Augier et al., 2016). mGluR2 receptors may mediate cognition within certain disease models by reducing cortical hyper-excitability (Gruber et al., 2010). Long term alcohol exposure similarly enhances cortical excitability (Nimitvilai et al., 2016; Cannady et al., 2018; Cannady et al., 2020; Hughes et al., 2020; Correas et al., 2021) suggesting that mGluR2 agonism may be a target that is capable of ameliorating differences in excitability.

Additional work is necessary to determine baseline cognitive differences between P rats and Wistars. de Falco and colleagues (2021) investigated cognitive flexibility between the rodent lines in an attentional set-shifting task. It was found that P rats show deficits in flexibility that may be attributable to urgency. It is therefore critical to refine our efforts to determine how proactive or reactive cognitive control changes over the course of a 2CAP session. For instance, Stock and colleagues (2023) found that, in humans, acute alcohol intoxication shifted cognitive control from reactive to proactive as a compensatory mechanism. Based on this as well as the results of de Falco (2021), it can be hypothesized that P rats’ proactive control may shift over the course of a session as urgency wanes, and they are forced to exert proactive control over behavior to counteract the effects of alcohol intoxication. This could be best measured with a block-design that shifts contingency in the middle of the session while the animals are already intoxicated. Additionally, if sucrose controls were utilized, we would expect to see more proactive control in the middle of the session but less compared with acute alcohol intoxication.

P rats and Wistars show differences in behavioral and electrophysiological correlates of cognitive control. P rats, that preferentially use reactive control, can readily maintain similar performance in our 2CAP task following a contingency change. Wistars, that preferentially use proactive control, are impaired following a contingency change. These data suggest that the dual mechanisms framework a potentially important and useful approach for investigating problematic alcohol use in preclinical rodent models that can then be directly translated to human, clinical populations.

## Notes

This work was supported by the following grants: T32-AA007462, RO1-AA029409, P60- AA007611. Additionally, the authors would like to acknowledge the comments of Cristine Czachowski, Nicholas Grahame, and Jonathan Brigman that were essential to shaping the present project.

### Competing Interest Statement

The authors have declared no competing interest.

